# Overglycosylation introduces local changes in triple helix alignment in collagen type I fibril structure

**DOI:** 10.1101/2025.11.09.687185

**Authors:** Luco Rutten, Judith Schaart, Liline A.S. Fermin, Jinge Xu, Marit de Beer, Robin H.M. van der Meijden, Karli R. Reiding, Anat Akiva, Nico Sommerdijk, Elena Macías-Sánchez

## Abstract

Collagen fibrils constitute the structural scaffold of bone, and their hierarchical organization is central to biomineralization. While hydroxylation and glycosylation of lysine residues are well-known collagen post-translational modifications, their structural consequences remain poorly understood. Here we show that excess of glycosylation of hydroxylysine residues leads to significant alterations in the local packing of collagen molecules within the fibrils. By prolonging the enzymatic modification window during helix folding through the use of cyclosporin A, an increase in double glycosylation was observed at specific sites. These overglycosylated residues were pinpointed by mass spectrometry, while cryogenic TEM revealed fibrils with reduced diameters and distinct displacements of defined sub-bands within the D-period, without altering the overall periodicity. By mapping the modified residues onto the quarter-staggered model, we have been able to correlate site-specific glycosylation with sub-band shifts, linking chemical modification to supramolecular order. These results provide molecular-level evidence that collagen glycosylation is an active determinant of fibril structure. Such insights not only advance fundamental understanding of collagen assembly but could also illuminate mechanisms underlying bone fragility disorders, including osteogenesis imperfecta, that feature altered glycosylation.

## Introduction

Bone tissue is a hybrid material with an extremely complex hierarchical structure that spans several orders of magnitude.^[1]^ It consists of an intricated extracellular matrix (ECM) mainly composed of collagen type I, the fibrillar protein that provides the structural scaffold, guiding the deposition of mineral.^[2]^ Because bone formation depends on the precise interplay between collagen fibril and mineral, the structural organization of the fibrils is essential for the mineralization process.^[3]^

Collagen is a triple helical protein containing three alpha-polypeptide chains with a repetitive amino acid motif of Gly-X-Y in which the X and Y position are frequently occupied by proline and hydroxyproline, respectively.^[4]^ Prior to the assembly of the triple helix, the polypeptides undergo post-translational modifications (PTMs), which are catalyzed by multiple enzymes. Prolyl-4-hydroxylase (P4H) catalyzes the hydroxylation of the proline on the Y position in the Gly-X-Y sequence of collagen.^[5]^ Lysyl hydroxylases 1-3 (LH1-3) are responsible for the 5-hydroxylation of lysine both in the triple helix, as well as in the terminal non-helical telopeptide segment.^[6]^ Hydroxylysine (Hyl) can be further modified by *O*-glycosylation to form galactosyl hydroxylysine (G-Hyl), which is catalyzed by the glycosyl transferase 25 domain 1 and 2 (*GLT25D1 and GLT25D2*).^[7]^ In a second glycosylation step, catalyzed by LH3, glucose can be coupled to the G-Hyl resulting in glucosylgalactosyl hydroxylysine (GG-Hyl).^[6]^

Before the formation of the triple helix, the ends of the polypeptides are bridged by disulfide bonds.^[8]^ Thereafter, the triple helix folds from the C-terminus to the N-terminus side. Once the triple helix is formed, the collagen molecules are exported to the extracellular space where they assemble in a quarter staggered array to form a fibril.^[9]^ This arrangement originates from the hydrophobic or electrostatic interactions of the outwards directed side chains of neighboring collagen molecules.^[10]^ These interactions are maximized at an axial displacement of 67 nm, forming zones of different molecular density that result in the so-called overlap (∼31 nm) and gap zones (∼36 nm),^[11]^ with an overall periodicity of ∼67 nm (the D-banding pattern).^[12]^ The D-band is further divided into multiple sub-bands, which are regions of aligned charged amino acids, resulting from the charge distribution in the collagen fibril.^[10, 13]^ Changes in the position of the sub-banding pattern can therefore be an indication of alterations in the spatial staggered arrangement of the collagen molecules.

The formation of covalent intra- and inter-molecular collagen crosslinks takes place outside the cell during fibril assembly, and it is mediated by the enzyme lysyl oxidase (LOX), which catalyzes the oxidation and deamination of lysine (Lys) to allysine, and of hydroxylysine (Hyl) to hydroxyallysine. These can undergo a condensation reaction with each other or can react with a Lys or Hyl in the triple helical domain of an adjacent collagen molecule to create bivalent crosslinks. These so-called immature crosslinks spontaneously evolve to form a variety of mature trivalent crosslinks .^[14]^ The most prominent trivalent crosslinks are lysylpyridinoline (LP) and hydroxylysylpyridinoline (HP).

Evidentially, collagen synthesis is a complex process with many interdependent steps. Glycosylation has been shown to have important functions related to collagen secretion,^[15]^ fibrillogenesis,^[16]^ organization of the ECM,^[17]^ crosslinking,^[18]^ and binding to non-collagenous proteins.^[19]^ Multiple studies have correlated the changes in fibril diameter with collagen modification.^[20]^ The decrease in fibril diameter as a result of an increase in glycosylation was previously reported by *in vitro* fibril assembly of soluble collagen with different degrees of glycosylation.^[20a, 20b]^ In these *in vitro* systems, the collagen was purified, so the role of non-collagenous proteins could be ruled out. The authors hypothesize that glycosylation modifies physicochemical properties, such as hydrophobicity, that could alter the fibril diameter. *In vitro* collagen glycosylation is also inversely correlated to accelerated fibrillogenesis.^[20c, 20d]^ Similarly, in mice were the Plod1 gene coding for LH1 was inactivated, the hydroxylation decreased and the fibril thickness increased.^[21]^ In addition, overglycosylated collagen is found in diseases with altered biomineralization, such as the “brittle bone disease” Osteogenesis Imperfecta (OI).^[22]^ Taken together, this suggests that changes in the pattern of glycosylation affect the structure of collagen and therefore could affect the mineralization process. However, the structural consequences of glycosylation are poorly understood.

Here, we test the hypothesis that increased glycosylation modifies fibril structure. Cyclosporin A (CsA) was added to mouse osteoblasts cell culture. CsA inhibits the peptidyl-prolyl cis-trans isomerization, which accelerates protein folding in living cells.^17^ Since proline isomerization is the rate limiting step in triple helix folding,^[23]^ a consequence of the inhibitory effect of CsA is a decrease in the folding rate of the triple helix.^[23]^ This results in an increase in the active time of the enzymes responsible for modifying the collagen polypeptide and a concomitant increase in the degree of glycosylation.^[23]^ By combining high performance liquid chromatography (HPLC)^[24]^ and mass spectrometry (MS) we determined the degree of glycosylation and the specific site of the overglycosylation on the amino acid sequence. These findings were correlated to the local changes in structure of the collagen fibrils determined by cryoTEM,^[25]^ suggesting that changes in the glycosylation pattern lead to significant changes in the collagen structure.

## Results & Discussion

### Overglycosylation of collagen from osteoblast under influence of CsA

In order to investigate the effect of the glycosylation on the collagen fibril structure, osteoblasts were cultured in the presence of CsA. To assess the viability of the osteoblasts and visualize the formation of collagen, the cultures were imaged with fluorescence microscopy at the beginning (4 days, **Fig. 1a-d**) and the end (28 days, **Fig. 1e,f**) of differentiation. Cultures were fixed and stained for collagen (CNA35-OG488),^[26]^ actin (phalloidin-Alex Fluor568) and nucleus (DAPI). After 4 days both cultures had produced collagen (**Fig. 1a,b**), but where the control culture had produced an intricated extracellular network. The CsA supplemented culture showed an increase of fluorescent signal around the nucleus of the cells, indicating collagen accumulation in the ER and a subsequent delayed collagen extrusion into the extracellular matrix (**Fig. 1c,d**). After 28 days of differentiation, both cultures showed a collagen network (**Fig. 1e,f**), although it was less dense in the CsA treated culture.

**Figure 1.**
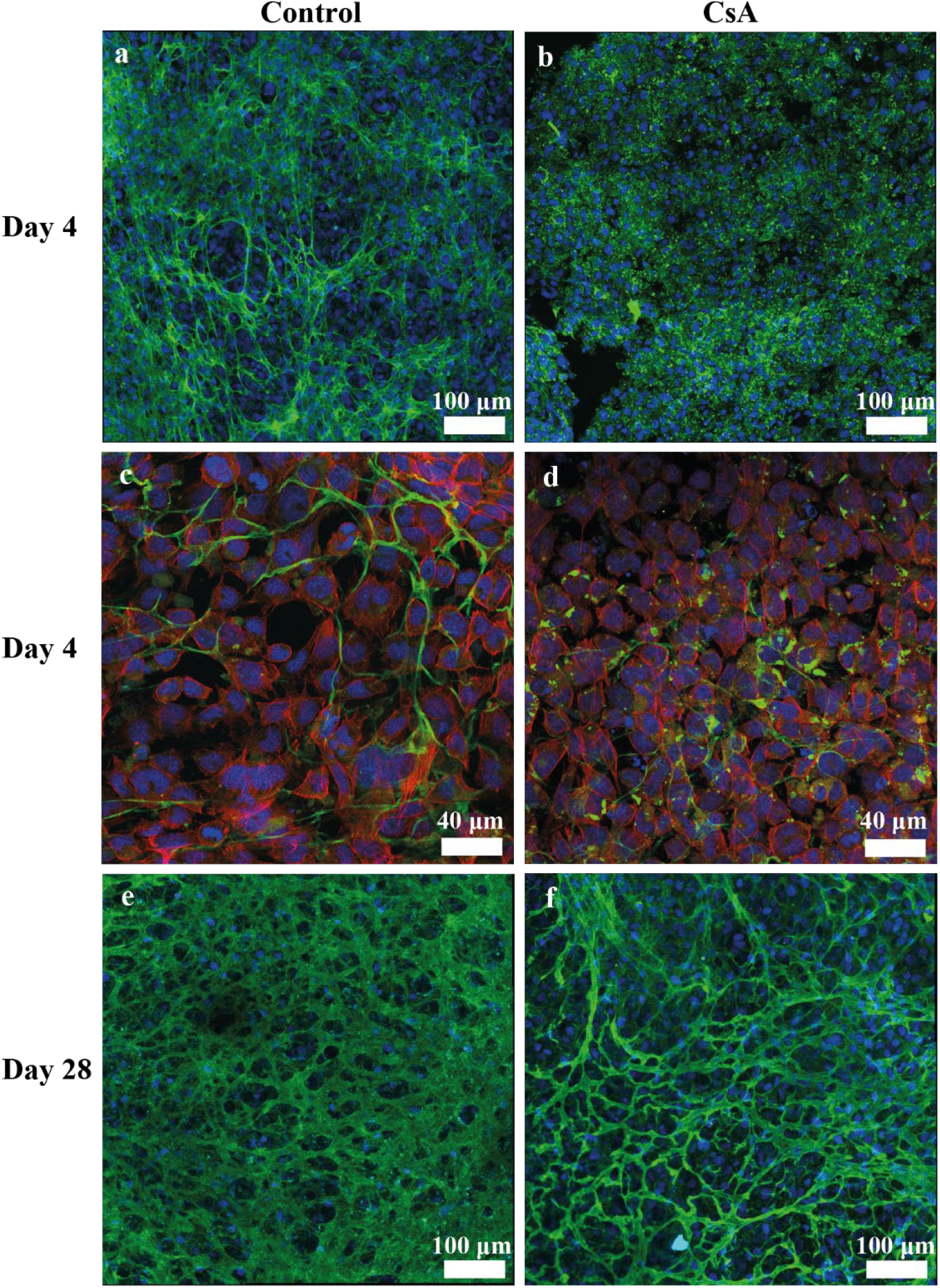
Fluorescence microscopy of collagen network formation for the control (a,c,e) and CsA supplemented (b,d,f) MLO-A5 cell cultures. Collagen (green), nucleus (blue) and actin (red). a,b) Collagen network and nucleus of the cells 4 days after differentiation. c,d) High magnification images of nucleus, actin and collagen 4 days after differentiation. e,f) Collagen network and nucleus after 28 days of differentiation.

This reduced density likely reflects altered secretion dynamics and increased intracellular retention of collagen. Intracellular accumulation of collagen could be related to an increase in glycosylation, as it has been suggested to play a role in the transport system of collagen out of the cell.^[15]^ On the other hand, CsA is known to have effects on cell metabolism, reducing osteoblast proliferation and altering bone protein synthesis (e.g. increased osteonectin expression),^[27]^ which could have affected the amount of collagen secreted. In any case, although the secreted amount may vary, our results showed that the amino acid composition of the synthetized collagen was similar to that of native collagen, indicating that slowing down collagen synthesis or transport does not affect the composition of the collagen (**Fig S1**).

### Biochemical characterization of isolated collagen

After isolation and purification of the collagen from the cell culture (**Supplementary text, Fig. S2,3**), HPLC with acid hydrolysis was used to quantify the total amount of hydroxylysine per collagen molecule (Hyl + GG-Hyl + G-Hyl). This acid treatment, performed at elevated temperature, hydrolyzes the peptide bond, but also hydrolyzes the glycosidic bonds attaching the carbohydrates.^[24]^ All G-Hyl and GG-Hyl products are therefore converted back to Hyl. The average total amount of Hyl per collagen molecule slightly increased from 54 to 59 by supplementing the cell culture with CsA (**Fig. 2b**), however this number was not statistically significant (p=0.08).

**Figure 2.**
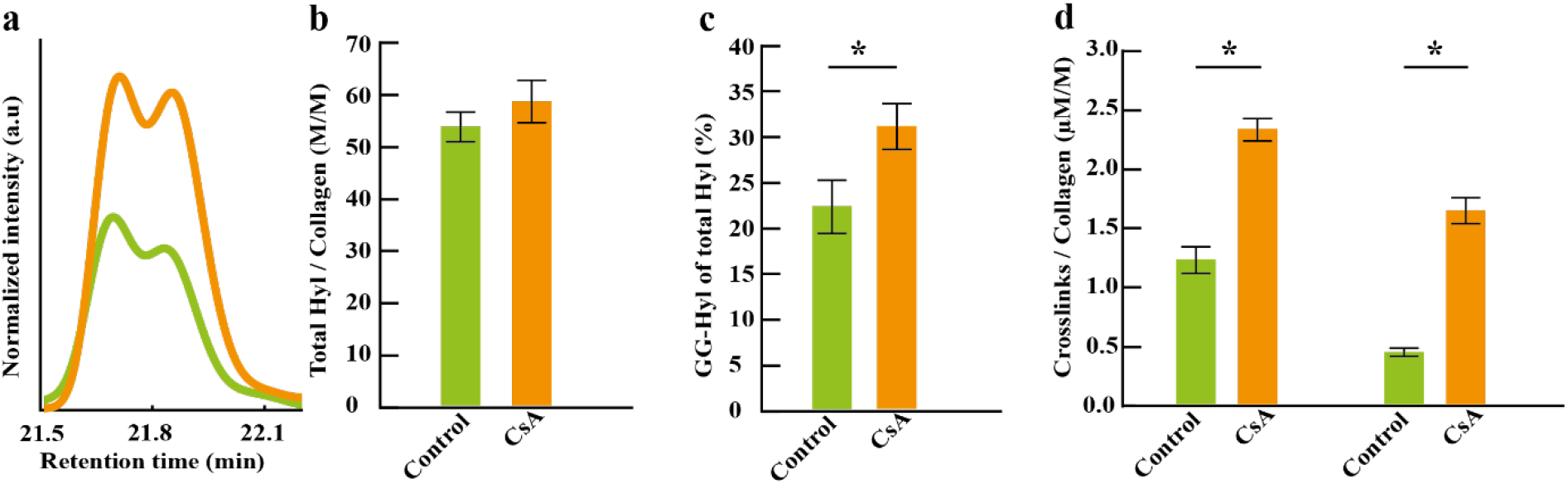
HPLC quantification of Hyl, GG-Hyl and mature crosslinks of control (green) and CsA treated (orange) collagen. a) Representative HPLC chromatogram of alkaline hydrolyzed isolated collagen at the specific retention time for GG-Hyl. The intensity is normalized to the Hyp peak. b) Average amount of hydroxylysine per collagen molecule. b) Average percentage of hydroxylysine that is glycosylated with glucosylgalactose (GG). d) Amount of hydroxylysyl pyridinoline (HP) and lysyl pyridinoline (LP) per collagen molecule. (*p=0.05)

Alkaline hydrolysis, which does not remove carbohydrates,^[24]^ was used to quantify the amount of Hyl that was glycosylated with glucosylgalactose (GG) (**Fig. 2a**). While the amount of hydroxylysine remained the same (**Fig. 2b**), the slowing down of collagen folding due to the CsA treatment led to a significant 50% increase of GG-Hyl (p=0.01) (**Fig 2c**). This is explained by the fact that GG-Hyl is the final product of the glycosylation process and accumulates as the enzymes have more time to interact with the collagen fibril. This accumulation was previously also observed in collagen produced by fibroblasts from OI patients^[22]^ and from peptidyl-prolyl cis-trans isomerase B (cyclophilin B) knock-out osteoblasts.^[20e]^ The amount of G-Hyl was below the detection limit in both the control and the CsA-treated sample.

Besides the degree of glycosylation, also the number of the most prominent mature crosslinks, hydroxylysyl pyridinoline (HP) and lysyl pyridinoline (LP), increased significantly (p=0.01) (**Fig. 2d**). An increase in the mature crosslinks stiffens the mechanical properties of collagen, but alterations in the crosslinking pattern could also influence the matrix mineralization.^[28]^ It was also suggested that double glycosylated immature crosslink do not have the tendency to mature to trivalent crosslinks.^[18]^ However, no glycosylated mature crosslinks were found, as these are very labile to acid treatment.^[29]^ Additionally, it has been shown previously that the absence of glycosylation decreases the number of trivalent crosslinks.^[30]^ Our results contradict this hypothesis, since both the glycosylation and mature crosslinks increased. Even though the factors controlling crosslink maturation are not understood yet, it is likely that the micro-environment as well as the position of the collagen molecules with respect to each other affect the maturation. The micro-environment could be influenced by the increased hydrophilicity of the carbohydrate groups and the steric effect of the bulky carbohydrate is shown to slightly alter the position of the molecule.^[31]^

The ratio between HP/LP (mean 1.99±0.7) is used as an indicator of bone strength.^[32]^ A higher HP/LP ratio correlates with increased stiffness of the bones.^[32]^ The HP/LP ratio decreased when comparing the CsA supplemented collagen (1.4) with control collagen (2.7).

The specific position and abundance of glycosylation are both tissue- and species-specific.^[6]^ Within a given tissue of a species, however, the abundance of glycosylation at a particular site is highly conserved. This high degree of specificity, together with the pathophysiological relevance on glycosylation, highlights the functional importance of site-specific glycosylation.^[33]^ Mapping these site-specific changes is essential, as the location and extent of overglycosylation could directly influence the supramolecular organization of the fibrils. Therefore, mass spectrometry was used to identify the specific amino acids in which additional glycosylation products were located. A key question is whether CsA-induced overglycosylation arises from increased modification at established glycosylation sites or from the introduction of new sites. Therefore, the identified overglycosylation sites were compared to the control as well as collagen type I from skin of various species, a tissue known for its high degree of glycosylation,^[18]^ since the glycosylation pattern of collagen type I in mouse bone remains poorly characterized.

Five lysine residues showed a significant increase in GG-Hyl: K_α1_509, K_α1_575, K_α1_731, K_α1_740, K_α1_770 (**Fig. 3a-e**). K_α1_509 and K_α1_731 (**Fig. 3a,c**) were already identified as glycosylation sites in a study on collagen type I from mouse skin.^[34]^ K_α1_575, and K_α1_770 (**Fig. 3b,e**) were previously identified as glycosylation sites on the collagen type I from the skin of deer, cow, goat and sheep^[35]^ and K_α1_770 also in collagen from human skin fibroblasts^[22]^. Interestingly, K_α1_740 was not identified as a potential glycosylation site in previous studies and shows a very low abundance (2%) in the collagen from the control (**Fig 3b**). However, when overglycosylation is induced, the relative abundance of GG-Hyl on K_α1_740 increases to 17%.

**Figure 3.**
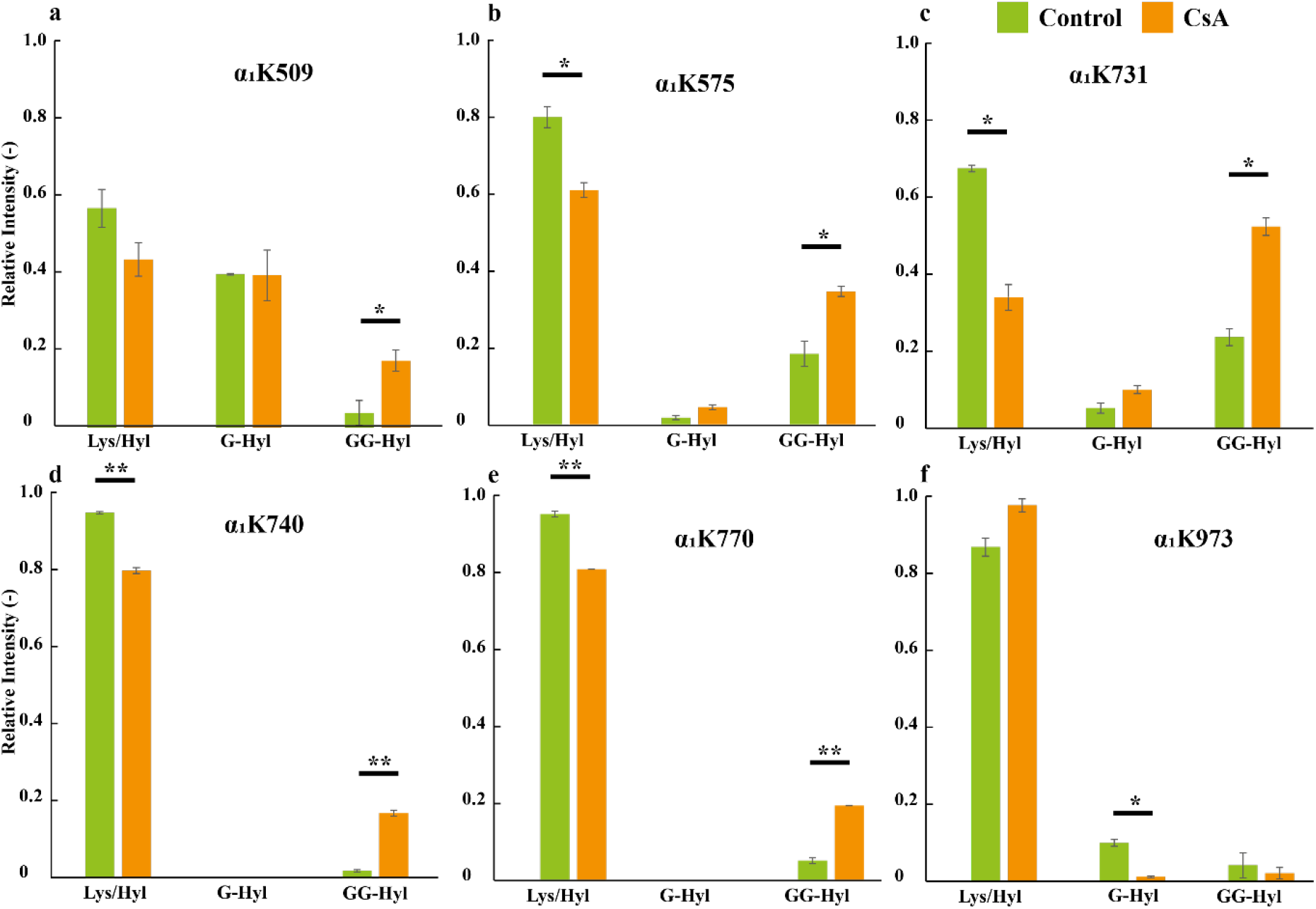
Identification of those lysine residues that showed significant differences in the hydroxylation and glycosylation pattern using mass spectrometry. Significant differences were found in a) α_1_K509, b) α_1_K575, c) α_1_K731, d) α_1_K740, e) α_1_K770 and f) α_1_K973 (* p<0.05, **p<0.005) between control (green, n=2) and CsA treated (orange, n=2) collagen. Graphs show the intensity of the peaks for lysine relative to hydroxylysine (Lys/Hyl), galgactosyl hydroxylysine (G-Hyl) and glucosyl galactosyl hydroxylysine (GG-Hyl).

Other known glycosylation sites (K_α1_254, K_α1_973, K_α2_1029), did show a relative abundance in glycosylation in both samples but did not show a significant increase (**Table S3**). Residues K_α2_660 and K_α2_852 previously reported as non-modified lysine or hydroxylation sites^[35]^ were found to be glycosylated in both samples, although they did not show a significant increase in CsA-treated samples (**Table S3**). Surprisingly, K_α2_973 showed a decrease in the relative abundance of G-Hyl when treated with CsA (**Fig. 3f**) but the relative abundance of GG-Hyl did not increase.

By slowing down the folding of the triple helix, the abundance of the glycosylation increased in specific lysines that were already glycosylated in the control sample. Since the number of residues is finite, the increase in the relative abundance of GG-Hyl in the CsA-treaded collagen entails a decrease in either Lys or Hyl. On another hand, the exact position of hydroxylation cannot be determined by mass spectrometry because it can occur in both proline and lysine residues, resulting in the same mass shift. That is why the decrease in the relative abundance of lysines after glycosylation is used to determine which residues were hydroxylated. In this regard, previous research on overglycosylated collagen produced by human fibroblasts from OI patients has also shown a decrease in Lys,^[22]^ indicating that the Lys residues are the ones being hydroxylated.

### Change in fibril structure investigated by cryoTEM

In several diseases that cause weaker bones (e.g. Osteogenesis Imperfecta and Ehlers-Danlos Syndrome), the collagen shows an increase in glycosylation^[36]^ together with a smaller fibril diameter^[37]^ compared to healthy bone collagen. To investigate if glycosylation could be responsible for this structural change in the fibril, isolated collagen fibrils from control and CsA-treated cell cultures were stained with uranyl acetate and visualized using cryoTEM (**Fig. 4a, Fig. S4**). The average diameter of the control fibrils was 34 nm (n=85), whereas that of the CsA-treated fibrils was 31 nm (n=117) (**Fig. 4b**). The total length of one D-band did not significantly change (**Fig. 4c**) showing that de CsA-treated collagen still assembles in the quarter-staggered arrangement.

**Figure 4.**
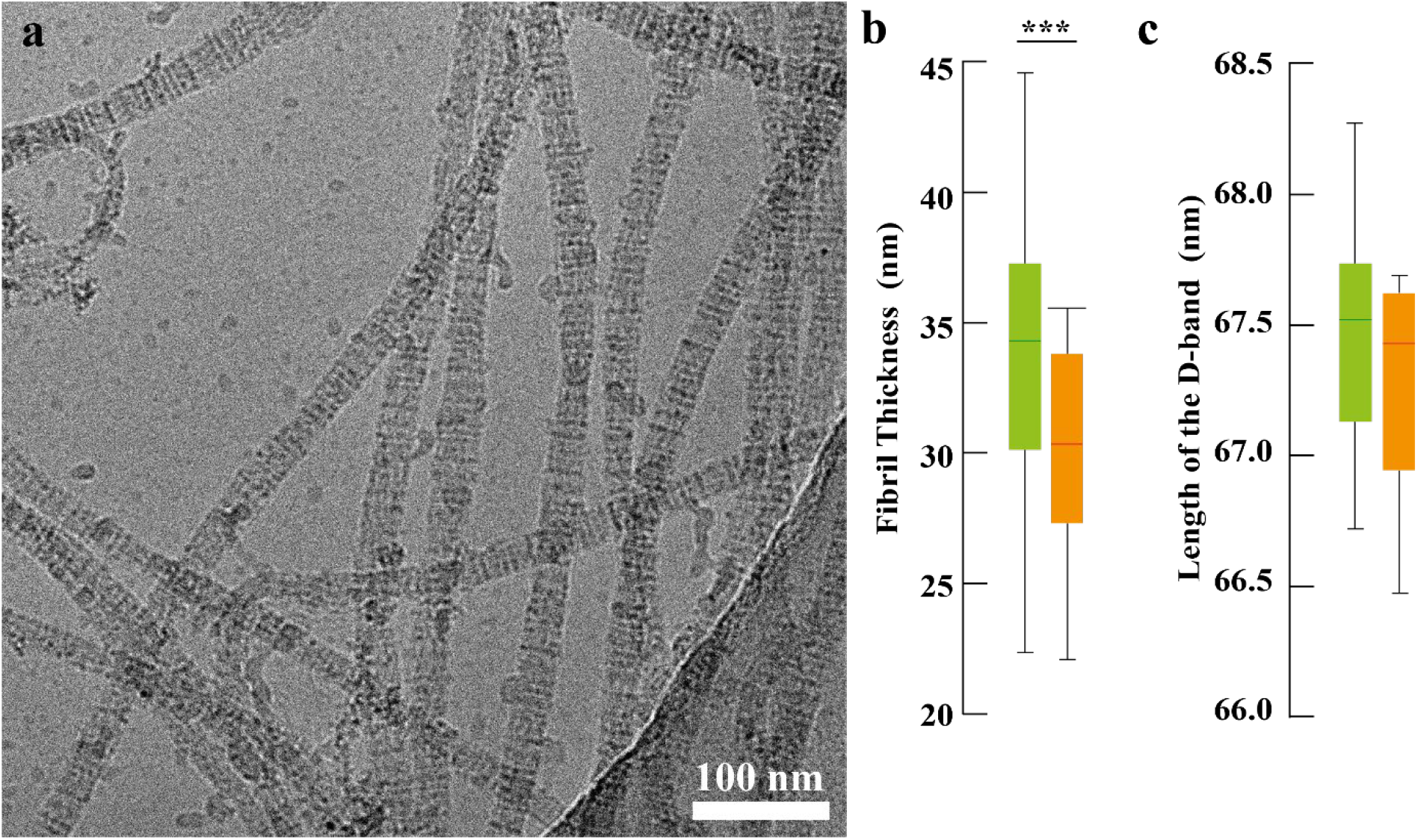
Analysis of collagen fibril structure. a) CryoTEM image of isolated uranyl acetate-stained collagen fibril from a cell culture treated with CsA. b) The average fibril thickness of completely assembled fibrils from control (green, n=85) and CsA treated samples (orange, n=115). c) Average length of the total D-band of control (green) and CsA-treated (orange) collagen fibrils. (***p<0.0005).

The sub-banding pattern was extracted from cryoTEM images of stained collagen fibrils (**Fig. S4**) isolated from cultures with similar passage number and known amounts of Hyl and GG-Hyl (**Table S2**). The difference between the control and treated collagen (**Fig. 5a**) and the average position of each sub-band (**Fig. 5b**) in one D-period (67 nm), are shown for the control (green) and CsA-treated (orange) fibrils (**Fig. 5b**). The position of the sub-bands of the control correspond well to the theoretical intensity profile (**Fig. 5b**) which is calculated based on the position of the charged (uranyl stained) amino acids in the 2D projected quarter staggered arrangement (**Fig. 5c**). However, in the CsA-treated sample four sub-bands (b1, a1, e1 & d) showed a significant change in their position when the glycosylation is increased by 70% from 7.2 to 12.5 GG-Hyl / collagen molecule (**Table S2**). The remaining sub-bands did not show any significant difference in the position.

**Figure 5.**
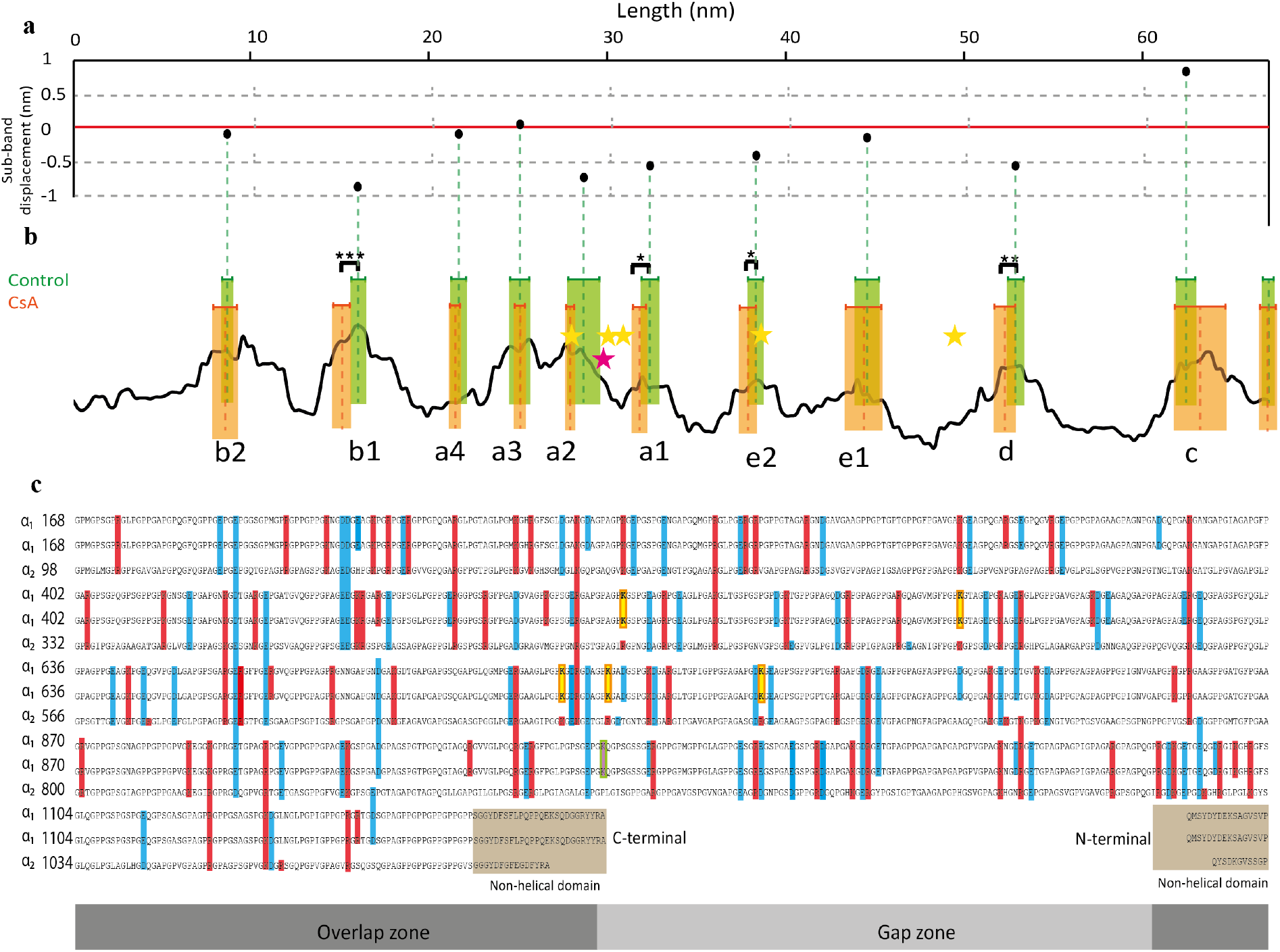
Analysis of collagen fibril sub-banding shifts. a) Black dots represent the relative displacement of the sub-bands in the CsA-treatment with respect to the control (red line) in one D-period (67 nm). b) Black line represents the theoretical intensity profile (simulated based on the mass of the amino acids) of a D-period of mouse collagen type I in a quarter staggered arrangement. The mass of uranium (238u) was added to all charged amino acids (highlighted in blue and red in c) and the sub-bands were named accordingly. The dotted line shows the position of the sub-bands of the control (green) and CsA (orange) samples with the error bars representing plus and minus one standard deviation. c) The amino acid sequence with the telopeptides of mouse collagen type I (Uniprot P11087 (α_1_), Q01149 (α_2_)) arranged in the quarter staggered arrangement showing one D-period (67nm).^[10]^ The positive (blue) and negative (red) charged amino acids are highlighted. The previously identified overglycosylation sites are highlighted in yellow (*p<0.05, **p<0.005, ***p<0.001).

Additionally, the mass spectrometry results enable to pinpoint the overglycosylated residues in de 2D arrangement (**Fig. 5c**) and spatially correlate them to the position in the sub-banding pattern (**Fig. 5b**). The broad peak between 28-37 nm in the displacement graph corresponds to four overglycosylation positions (K_α1_731, K_α1_740, K_α1_509 and K_α1_770), where the relative abundance of GG-Hyl for K_α1_731, K_α1_740, K_α1_509 and K_α1_770 increases from 25% to 54%, 2% to 18%, 4% to 17% and 5% to 19%, respectively (**Fig. 3a-c,e**). Glycosylation site K_α1_575, (**Fig. 3d)**, is in close proximity to the d sub-band which also shows significant displacement. The displacement in the b1 sub-band cannot be readily correlated to a specific overglycosylated residue. However, K_α2_852 and K_α1_923, which are part of the b1 sub-band, do show a non-significant increase in relative abundance of GG-Hyl (74% to 89%, p=0.11) and G-Hyl (37% to 70%, p=0.06), respectively (**Table S3**).

The correlation of the specific glycosylation sites with the position of the sub-bands shows a relationship between glycosylation and fibril structure. Since glycosylation increases the hydrophilicity of the Hyl but also increases the bulkiness of the side group, both factors may impact the assembly of collagen molecules. The D-spacing is mainly determined by ionic and hydrophobic intra-molecular interactions and is fixed in place by crosslinking of specific (hydroxy)lysine in the telopeptide and on the triple helix. If glycosylation is introducing variability in the inter-residue spacing, this could also explain the changes observed in the sub-bands while observing no changes in the total length of the D-band. A computational study based on X-ray diffraction patterns of hydrated collagen showed that including not-uniform axial pitch in the triple helix improves the fit of the calculated model to the experimental data from rat tail collagen.^[38]^

Molecular simulations of glycosylated collagen have shown that interactions of the carbohydrates could alter the protein backbone conformation and the local structure of the water molecules.^[39]^ In addition, glycosylation can change the local solvation around the glycosylation sites, which is an important parameter in collagen assembly.^[40]^ An FFT analysis on the position of glycosylated residues in a collagen molecule showed a relationship between the linear distribution of the glycosylated Hyl, and the D-spacing observed in collagen fibril.^[35]^ This means that the glycosylated moieties can only interact with each other (to generate e.g. cross-links) when the fibril is properly assembled. Our experimental observation suggests a structural/functional role of glycosylation in collagen assembly and supports the hypothesis that increased glycosylation could alter the fibril self-assembly.

Our results showed that collagen molecules with increased level of glycosylation were still able to assemble in a quarter staggered array with a periodicity of ∼67 nm. However, CsA-treated fibrils had a smaller diameter (p=3.4·10^-5^). The relation between fibril diameter and glycosylation was also previously shown and attributed to steric interactions.^[41]^ We confirm the morphological changes by showing alteration in the sub-banding pattern. The location of the overglycosylation in the axial projection of the quarter staggered arranged fibril suggests that local changes in the axial pitch as a result of overglycosylation could explain the shifts in the sub-bands.

Although the role of collagen glycosylation in health and disease is not completely understood, changes in collagen glycosylation have a severe impact on the structure/function relationship and thereby alter the properties of the ECM.^[33]^ In the case of the biomineralization of bone the changes in the self-assembly could affect the packing of the collagen.^[42]^ Since the collagen confinement facilitates the growth of mineral,^[3]^ this change could alter the mineral growth and the properties of the bone. Therefore, the structural changes in the collagen fibrils due to overglycosylation may contribute to more fragile bones as observed in certain disease which are characterized by both glycosylation and bone deficiencies such as Osteogenesis Imperfecta^[22, 43]^, osteopenia^[44]^ and Ehlers-Dalos syndrome^[45]^.

## Conclusion

We demonstrate that overglycosylation of collagen introduces localized misalignment in triple helix packing, detectable as shifts in sub-banding within the fibril D-period. By correlating specific lysine glycosylation sites with nanoscale structural changes, we establish glycosylation as a molecular regulator of supramolecular collagen order. This chemical–structural link provides a new framework for understanding how subtle post-translational modifications can propagate into tissue-level defects in biomineralization and bone mechanics.

## Materials & Methods

### Cell culture

Mouse osteoblasts MLO-A5^[46]^ were cultured in proliferation medium containing α-MEM supplemented with 5% bovine calf serum (BCS) and 5% fetal calf serum (FCS). For differentiation the αMEM was supplemented with 10% FCS, 100 µg/mL ascorbic acid and Cyclosporin A (CsA) dissolved in ethanol or pure ethanol for the control. This resulted in a final CsA concentration of 5 µM. The medium was exchanged every 2-3 days.

### Fluorescent microscopy

Cell culture samples on cover slips were washed with PBS, and then fixated with 4% paraformaldehyde in PBS for 15 minutes. The PFA was diluted to 1% by addition of PBS. The samples were stored at 4°C until staining. Before staining, the coverslips in the well plate were washed 3 times with PBS. Staining solution was prepared with CNA-OG488 (collagen, TU/e) at a dilution of 1:50, Phalloidin Alexa Fluor 568 (actin, Thermo Fisher Scientific) 1:250 and DAPI (nucleus, Thermo Fisher Scientific) 1:500 in PBS. Drops of 15μL staining solution were placed on parafilm and the coverslips were placed on top. Coverslips were left in the dark for 1hr to stain. Coverslips were then transferred back into a well plate to wash twice with PBS. The coverslips were taken out of the well plate and the back of the coverslip was dried. They were then placed on a filter paper, and a drop of polyvinyl alcohol (Mowiol) was added on top. A microscope slide was then gently placed on top of the coverslips, and the microscope slide was then transferred to a container to cure overnight at room temperature. The Zeiss LSM 900 was used to image the samples, and ImageJ was used for analysis.

### Collagen isolation

After 5 weeks of differentiation the collagen was isolated using a three-step isolation protocol: 1) freeze/thawing procedure, 2) detergent clean-up and 3) enzymatic digestion. All the working buffers and washing solutions were prewarmed to 37°C before use.

#### 1) Freeze/thawing

To isolate the collagen from the cell culture, the culture was washed twice with PBS and frozen at -80°C. The frozen cell culture was thawed at room temperature and washed twice with PBS and once with Milli-Q.

### 2) Detergent clean-up

The cells were washed twice with PBS whereafter the extraction buffer was added (20mM NH_4_OH, 0.5% Triton X in PBS, pH 8). Cell culture with extraction buffer was incubated at 37°C for 10 min. The sample was transferred to a centrifuge tube and diluted twice with PBS. After overnight rotation at 4°C, the extraction buffer was removed, and the residue was washed twice with PBS. The sample was incubated with a DNase buffer (10μg/mL DNase in 100mM Tris-HCl buffer with 25mM MgCl_2_, 1mM CaCl_2_) at 37℃ for 30min. The sample was washed by DNase working buffer (100mM Tris-HCl buffer with 25mM MgCl_2_, 1mM CaCl_2_), PBS and twice with Milli-Q respectively.

#### 3) Enzymatic digestion

The sample was incubated in α-chymotrypsin buffer (0.05mg/mL α-chymotrypsin in 100mM Tris-HCl buffer with 10mM CaCl_2_) at 37°C for 20 hours after being 2 times washed by enzyme working buffer (10mM CaCl_2_ in 100mM Tris-HCl buffer). The enzymatic digested sample was washed by Milli-Q, two times PBS and Milli-Q respectively. 0.25%(v/v) Tween-20 was added to sample and removed after incubation at 37°C for 20 hrs. The sample was washed twice with PBS and twice with Milli-Q. The purified collagen sample was stored in Milli-Q water at 4°C until further research.

### Hydrolysis and derivatization

HPLC analysis was preformed according to the protocol of Bank et al^[47]^ with some modification. In short, the decellularized collagen matrix was lyophilized and hydrolyzed for 24h at 110 °C using either 6M HCl or 2M NaOH. For the acidic hydrolysis the HCl was evaporated overnight using a vacuum desiccator. The remaining hydrolysate was dissolved Milli-Q and diluted in 0.1 M borate buffer. After hydrolyzed with NaOH the solution was neutralized with an equivalent amount of 2 M HCl and subsequently diluted in 0.1 M borate buffer. Homoarginine was added as internal standard to the sample. Afterwards the sample was derivatized with 6 mM fluorenylmethyloxycarbonyl (FMOC) and prepared for HPLC analysis according to the previous established protocol.

### HPLC analysis

The HPLC system (Prominence, Shimadzu Scientific Instruments) equipped with RF-10AXL detector and reversed phase column (TSKgel ODS-80Tm, 4.6 mm ID × 15 cm L, 5 µm pore size). A solution containing 22.3 mM citric acid, 5 mM tetramethylammonium chloride and 0.01 (m/v) sodium azide was adjusted to pH 2.85 using a solution containing 19 mM tri-sodium citrate dihydrate, 5 mM tetramethylammonium choride and 0.01 (m/v) sodium azide to generate Eluent A. Eluent A was filtered through 0.2 μm pore size Whatman^®^ membrane. HPLC-grade acetonitrile was used as Eluent B. Sample was ejected on the column (40 °C) and the fluorescent signals were detected at 254nm (excitation) and 600nm (emission) at a total flowrate of 1.4 ml/min.

### Cross-links analysis protocol

For each sample, the chromatographic gradient of Eluent X, Y and Z is described in Table 3 below. The flow rate was maintained at 1 mL/min throughout each analysis. Derivatized internal standard samples and collagen samples (25-100 μL, depending on samples) were injected to the HPLC column at 40°C. The fluorescent signals were detected at 295nm (Excitation) and 400nm (Emission) from 0-20 min; at 328 nm (Excitation) and 378 nm (Emission) for 20-30 min.

**Table 2.** HPLC elution gradient for derivatized amino acid analysis.

| Time (min) | Eluent A (%) | Eluent B (%) |
| --- | --- | --- |
| 0 | 75 | 25 |
| 11.5 | 60 | 40 |
| 13 | 60 | 40 |
| 13.1 | 64 | 36 |
| 18 | 62 | 38 |
| 25 | 30 | 70 |
| 30 | 25 | 75 |
| 32 | 25 | 75 |
| 32.1 | 75 | 25 |
| 40 | 75 | 25 |

**Table 3.** HPLC elution gradient for cross-links analysis.

| Time (min) | Eluent X (%) | Eluent Y (%) | Eluent Z (%) | Flow rate (mL/min) |
| --- | --- | --- | --- | --- |
| 0 | 100 | 0 | 0 | 1.0 |
| 17 | 0 | 100 | 0 | 1.0 |
| 30 | 0 | 0 | 100 | 1.0 |
| 40 | 100 | 0 | 0 | 1.0 |
| 50 | 100 | 0 | 0 | 1.0 |

Cross-link Eluent X consisted of 0.12% (v/v) heptafluorobutyric acid (HFBA) in 24% (v/v) methanol in Milli-Q. 0.07% (v/v) HFBA in 40% (v/v) methanol in Milli-Q water was used in Eluent Y. Eluent Z contained 0.15% (v/v) HFBA in 75% (v/v) acetonitrile in Milli-Q. The cross-link (CL) sample buffer was 10% (v/v) acetonitrile in Milli-Q water containing 1% (v/v) HFBA. Eluent X and Y were filtered through 0.2μm pore size Whatman^®^ membrane. All eluents were prepared by HPLC-grade reagents.

The CL internal standard containing 1.5 μM pyridoxine (VB6), 1.5 μM pentosidine (PEN), 5.8μg/mL pyridinoline (PYD) and 3.06μg/mL deoxypyridinoline (DPD) (PYD/DPD HPLC calibrator, Quidel) were made in the CL sample buffer. 25 μL of collagen sample (in Milli-Q) was added to 125 μL CL sample buffer containing 1.5 μM pyridoxine.

#### Cryo-Transmission Electron Microscopy

Gold R2/2 Quantifoil grids were plasma cleaned for 40 seconds using a Cressington 208 coater. The collagen suspension was applied on a grid and subsequently the excess liquid was blotted away. Thereafter, collagen was stained with 2% uranyl acetate for 2 minutes. The grids were vitrified using a FEI Vitrobot™. A JEOL JEM-2100 equipped with a LaB_6_ filament and operated at 200 kV was used for imaging. The software Serial EM was used to acquire images in low-dose mode.

#### Mass spectrometry

Samples were reduced, alkylated and digested by trypsin as described previously.^[48]^ An approximate 200 ng of each digested sample was analyzed by LC-MS/MS, i.e., reversed phase chromatography on an Ultimate 3000 HPLC system (Thermo Scientific, Bremen, Germany) coupled to an Orbitrap Exploris 480 mass spectrometer (Thermo Scientific, Bremen, Germany). For the chromatography, we used a 2 cm trap column (in-house packed ReproSil-Pur C18-AQ, 3 μm; Dr. Maisch GmbH, Ammerbuch-Entringen, Germany) followed by a 50 cm 50 µm inner diameter analytical column (in-house packed Poroshell 120 EC-C18, 2.7 μm; Agilent Technologies, Amstelveen, The Netherlands). The separation was performed for 60 min with mobile phases of 0.1% formic acid in water (buffer A) and 0.1% formic acid in acetonitrile (buffer B). The gradient started with 9% B, going up to 13% B in 1 min, up to 44% B across 40 min, then up to 99% B in 1 min, held at 1 min, followed by a washing step of 10 min at 9% B. The flow rate was set at 300 nL/min.

Samples were ionized at a spray voltage of 2000 V, with the ion transfer tube heated to 275 °C. MS1 spectra were acquired for *m/z* 350-2000 at a resolution of 120,000, to reach a “standard” AGC target within 50 ms. Within a cycle time of 3 s, precursors, starting with the highest intensity and a minimum intensity of 50,000, charge 2-8+, were isolated for fragmentation to a “standard” AGC target and maximum accumulation time of 50 ms. Selection for MS2 led to an exclusion time of 30 s. Precursor fragmentation was performed by HCD at a normalized collision energy of 29%, followed by measurement in the Orbitrap analyzer at a resolution of 60,000 and with a scan range of *m/z* 120-4000.

Database searches were performed with Byonic (v5.7.11, Protein Metrics) using as small, dedicated database containing mouse CO1A1 (P11087) and CO1A2 (Q01149) in addition to common contaminants and reversed sequences. The searches allowed semi-specific tryptic peptides up to 5 miscleavages, a precursor tolerance of 6 ppm and a fragment tolerance of 20 ppm. The common modification limit was set to 4 and that of the rare modification to 1. Amongst the modifications, C carbamidomethylation was set to fixed, M/W oxidation to common 2, N-term Q/E pyroglutamic acid formation to common 1, S/T/Y phosphorylation to common 1 and protein N-term acetylation to common 1. Next to this, 279 *N*-glycans were included as rare 1, 5 *O*-glycans as rare 1, as well as W *C*-mannosylation as rare 1. To analyze collagen modifications specifically, we included as common 2 the formation of K hydroxylysine (+15.99), P hydroxyproline (+15.99), K galactosylhydroxylysine (+178.04) and K glucosylgalactosylhydroxylysine (+340.10). Search results were curated to have a score of ≥ 150 and |log prob| ≥ 1.5, and limited to have a false discovery rate of <1% on basis of the reverse detections.

To analyze the areas of modified and miscleaved peptides, we made use of Skyline (64-bit, v25.1.0.142). Byonic-reported peptides, including modifications and miscleavages, were loaded into Skyline and used to search and integrate the raw data files. Skyline integrations were then manually curated to 1) adhere to the retention times reported by the Byonic search (± 2 min), 2) to have a dot-product (idotp) between theoretical and observed isotopic distributions of at least 0.8 (in the best replicate), and 3) to have an error between theoretical and observed *m/z* values of at most 6 ppm. After area integration, charge states and miscleavages were summed to arrive at composite values for lysine modification.

*Statistics:* The means between two groups was compared by using t-tests (Analysis ToolPak, Excel). For the overglycosylation a one-side t-test with equal variance was used and for the position of the sub-bands a two sides t-test with equal variance.

## Acknowledgements

LR, JS, MdB, LF, JX, RvdM, AA, and NS were supported by the European Research Council (ERC) Advanced Investigator grant (H2020-ERC-2017-ADV-788982-COLMIN). EMS was supported by the Ramón y Cajal Program (RYC2023-045512-I) funded by MCIN/AEI/10.13039/501100011033 and FSE+ and the project PID2022-141993NA-I00 funded by MICIU/AEI/10.13039/501100011033 and ERDF/UE.

