## Supplementary material for "Overglycosylation introduces local changes in triple helix alignment in collagen type I fibril structure"

### Supporting information

#### Collagen purification

The isolation of collagen fibrils from all the other components in the cell culture, such as proteins, DNA, cell debris, lipids, etc. is a crucial step. The most frequently used method to isolate collagen from a cell culture is washing with a detergent followed by lyophilization.[49] However, chemical characterization of the obtained collagen with Raman spectroscopy showed this method does not result in high purity collagen (**Fig. S2a**). The Pro (~855 $cm^{-1}$) and Hyp (~875 $cm^{-1}$) peaks are low compared to the amide peaks (~1245 $cm^{-1}$, ~1660 $cm^{-1}$), indicating contamination with other proteins. Besides, the peak at ~1360 $cm^{-1}$ is indicating contamination (CH vibration of DNA/RNA).[50]

We therefore applied a optimized isolation and purification protocol to the collagen synthetized after 4 weeks of differentiation.[51] The protocol consists of three steps: 1) freeze/thaw cycle, 2) detergent washing and 3) enzymatic digestion. The freeze/thaw step ruptures the cell membranes which are washed away by the detergent (**Fig. S2b**). Enzymatic digestion with α-chymotrypsin removes the non-collagenous proteins (**Fig. S2c**). Raman spectroscopy confirmed the isolated product had chemical characteristics similar to that of reference collagen, implying that the enzymatic digestion step is crucial for obtaining high quality collagen (**Fig. S2d**).

Subsequently the collagen was lyophilized and hydrolyzed using acidic hydrolysis to analyze the amino acid composition of the collagen using high performance liquid chromatography (HPLC) (**Fig. S3**). Since the hydroxylation of proline is a specific modification of collagen, the degree of purity of collagen can be estimated by determining the Hyp/Pro ratio. All Pro residues at the Y-position in the [Gly-X-Y] sequence are 4-hydroxylated.[6] A decrease in the Hyp/Pro ratio implies higher Pro content, meaning the sample contains proteinaceous impurities. The average Hyp/Pro ratio of the purified collagen (0.87±0.02) compared the Hyp/Pro ratio of demineralized bone (0.81) or rat tail collagen (0.80) were higher (**Table S1**), indicating the purity of collagen after the enzymatic purification was excellent.

Ratios of the relative contribution of the other amino acids were comparable to collagen type I reference (**Fig. S3a,b**). Furthermore, the composition of control and CsA treated collagen was similar, confirming that CsA treatment did not affect the amino acid sequence **(Fig. S3c,d, Fig S1**). This isolation method results in pure pristine collagen and confirms the validity of using an osteoblast cell culture for the investigation of the self-assembly process.

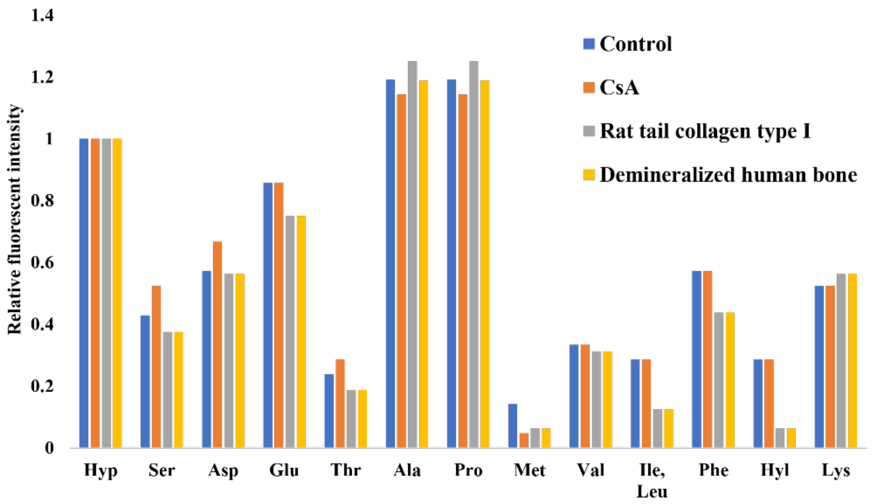


**Figure S1** Relative fluorescent intensity of the amino acids using HPLC

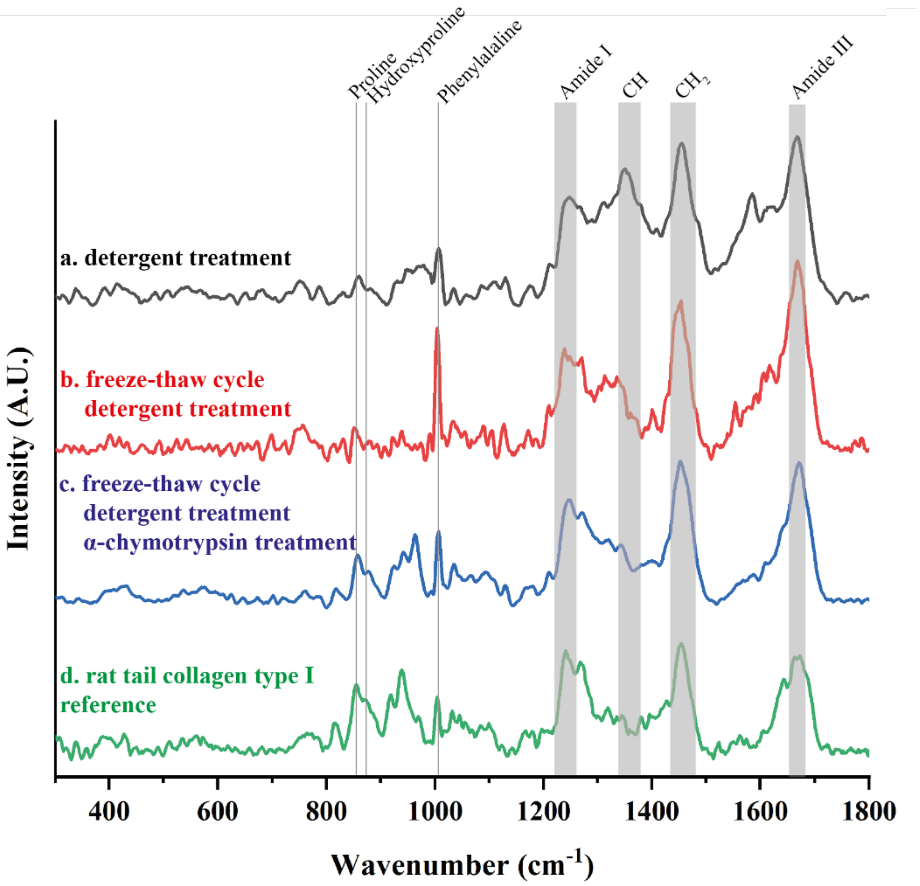


**Figure S2.** Raman spectra of collagen obtained by different purification protocols compared to the spectrum of commercial type I rat tail collagen. Characteristic peaks for collagen are indicated. a) Detergent treatment (black). b) Freeze-thaw cycle and subsequent detergent treatment (red). c) Freeze-thaw cycle, detergent treatment, enzymatic treatment with α-chymotrypsin and subsequent detergent wash (blue). d) Commercially available rat tail type I collagen (green) (C7661, Sigma Aldrich).

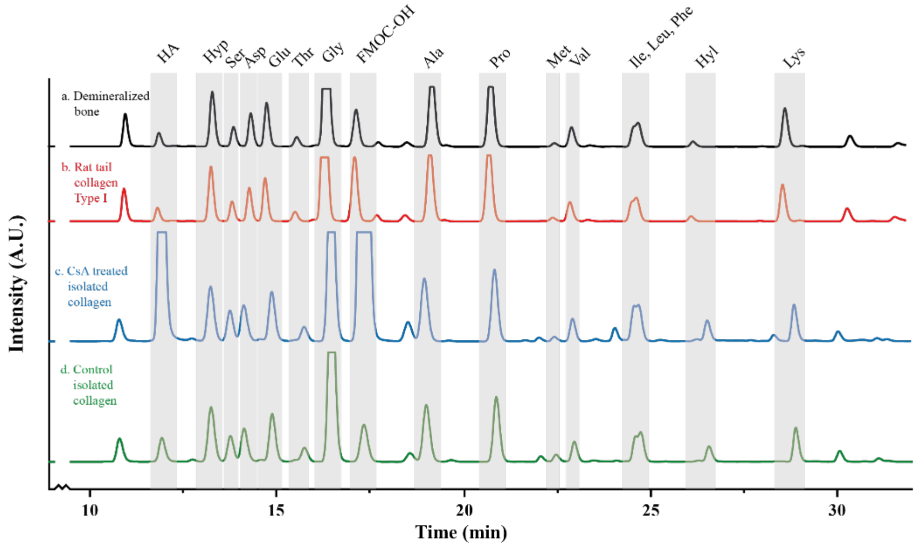


**Figure S3.** Chromatograms of different collagen samples acquired with HPLC after acid hydrolysis. a) Demineralized bone. b) Commercially available rat tail collagen type I. c) Collagen isolated from an MLO-A5 culture treated with CsA. d) Collagen isolated from an MLO-A5 culture under standard conditions. (HA: homoarginine (internal standard), Hyp: hydroxyproline, Ser: serine, Asp: Aspartic acid, Glu: Glutamic acid, Thr: threonine, Gly: glycine, FMOC-OH: fluorenylmethyloxycarbonyl hydroxyl, Ala: Alanine, Pro: proline, Met: methionine, Val: valine, Ile: isoleucine, Leu: leucine, Phe: phenylalanine, Hyl: hydroxylysine, Lys: lysine).

**Table S1**. Hydroxyproline to proline (Hyp/Pro) ratio of isolated collagen compared to collagen references.

| | **Isolated collagen** | **Demineralized bone** | **Rat tail collagen type I** |
|---|---|---|---|
| Hyp/Pro ratio | 0.87±0.02 | 0.81 | 0.80 |

**Table S2** Number of Hyl and GG-Hyl per collagen molecule of the fibrils for which the sub-banding pattern was extracted for structural analysis.

| Sample | Total Hyl / Collagen | % GG-Hyl of total Hyl | GG-Hyl / Collagen |
|---|---|---|---|
| Control | 51.8 | 7.2 | 3.7 |
| CsA | 56.5 | 12.5 | 7.1 |

**Table S3.** Position and relative abundance of overglycosylation on specific lysine

| Number of Lysine | | Control | | | | CsA | | | | t-test | Ref | | |
|---|---|---|---|---|---|---|---|---|---|---|---|---|---|
| Begin protein | Begin triple helix | Lys | Hyl | G-Hyl | GG-Hyl | Lys | Hyl | G-Hyl | GG-Hyl | p-value | a | b | c |
| α1-K254 | α1-K87 | 0.01 | 0.22 | 0.00 | 0.77 | 0.01 | 0.39 | 0.00 | 0.61 | | GG | GG | GG |
| α1-K341 | α1-K174 | 0.12 | 0.88 | 0.00 | 0.00 | 0.08 | 0.92 | 0.00 | 0.00 | | GG | GG | GG |
| α1-K509 | α1-K342 | - | 0.57 | 0.40 | 0.04 | 0.00 | 0.43 | 0.39 | 0.17 | 0.04 | G | Lys | |
| α1-K541 | α1-K374 | - | 0.89 | 0.11 | | - | 0.99 | 0.01 | | | Lys | Lys | |
| α1-K575 | α1-K408 | - | 0.80 | 0.02 | 0.18 | - | 0.61 | 0.04 | 0.35 | 0.02 | Hyl | GG | |
| α1-K731 | α1-K564 | - | 0.70 | 0.06 | 0.25 | - | 0.35 | 0.10 | 0.54 | 0.01 | G | Hyl | |
| α1-K740 | α1-K573 | - | 0.98 | 0.00 | 0.02 | - | 0.82 | 0.00 | 0.18 | 0.00 | Lys | Lys | |
| α1-K770 | α1-K603 | - | 0.95 | 0.00 | 0.05 | - | 0.81 | 0.00 | 0.19 | 0.00 | Lys | G | GG |
| α1-K923 | α1-K756 | - | 0.63 | 0.37 | 0.00 | - | 0.30 | 0.70 | 0.00 | | Lys | G | |
| α1-K973 | α1-K806 | - | 0.86 | 0.10 | 0.04 | - | 0.97 | 0.01 | 0.02 | 0.01 | Lys | G | |
| α2-K183 | α2-K87 | - | 1.00 | | | - | 1.00 | | | | Hyl | GG | GG |
| α2-K660 | α2-K564 | - | 0.52 | 0.22 | 0.26 | - | 0.13 | 0.43 | 0.44 | | Lys | Hyl | |
| α2-K747 | α2-K651 | x | x | x | x | X | x | x | x | x | Hyl | | |
| α2-K852 | α2-K756 | - | 0.26 | | 0.74 | - | 0.11 | | 0.89 | | Lys | Lys | |
| α2-K980 | α2-K884 | - | 1.00 | | | | 1.00 | | | | Lys | Hyl | |
| α2-K1029 | α2-K933 | - | 0.96 | | 0.04 | | 0.94 | | 0.06 | | Hyl | G | |
| α2-K1070 | α2-K974 | 0.93 | 0.01 | 0.01 | 0.05 | 0.96 | 0.03 | 0.00 | 0.01 | 0.02 | Lys | Lys | |

[a] 98

[b] D. R. Visser, T. S. Loo, G. E. Norris, D. A. D. Parry, *Journal of Structural Biology* **2023**, *215*, 107938
[c] Y. Taga, M. Kusubata, K. Ogawa-Goto, S. Hattori, *Journal of Proteome Research* **2013**, *12*, 2225-2232.

### Uranyl staining

Due to the partial dissociation of the uranyl acetate in aqueous solution, multiple uranium-containing complexes, anionic as well as cationic, are produced.[52] Therefore, the contrast is based on the reaction of these complexes with positive ($=NH_2^+$ and $-NH_3^+$ of Arg and Lys) and negative ($COO^-$ of Asp and Glu) groups of amino acids. The amount of staining is thus depending on the reactivity of the charged groups.[53] UA has the advantage that is has a small grain size (4-5 Å) compared to other heavy metal stainings.[54] This is necessary to obtain the resolution for visualizing the subtle changes in the sub-banding of the collagen fibrils.[10] Since the local quality of the staining also influences the intensity of the sub-bands, only the position of the bands and not the intensity was considered. Lys hydroxylation and glycosylation could result in a slight decrease in the reactivity of the amino groups. However, as these modifications do not react with the amino group, the charge of the side groups is not expected to change. HPLC analysis has shown that the amino acid sequence of the overglycosylated fibrils remains unaltered, and therefore the sub-banding pattern of both, control and overglycosylated fibrils, results from the interaction of the UA with the same amino acids.

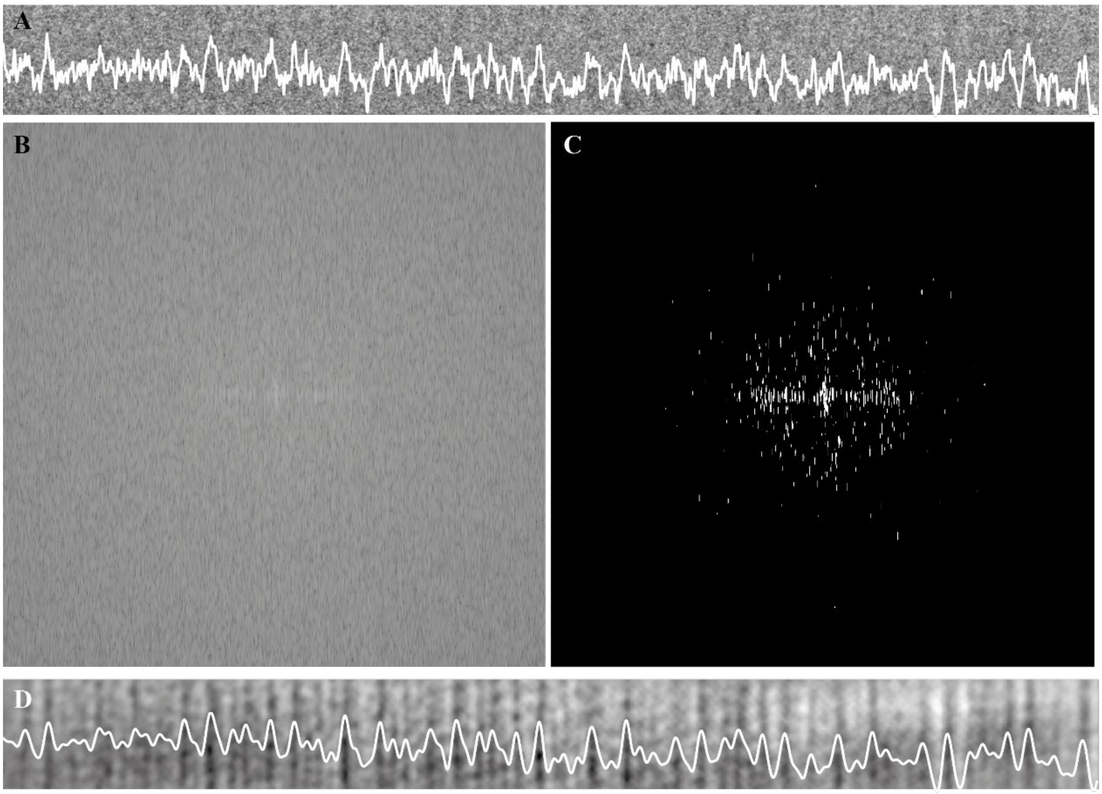


**Figure S4.** Extraction of the intensity profile. a) Th line shows the intensity profile of a cropped area from a cryoTEM image on a uranyl acetate stained fibril. Each darker vertical line, corresponding to a peak in the intensity plot, indicates one sub-band. To extract the precise position of each sub-band, the fast Fourier transformation (FFT) of the cropped area was taken (**Fig. 5b**) and a triangle threshold applied in Fiji (**Fig. 5c**). The inverse FFT shows an enhanced sub-banding pattern without altering the position of the peaks (**Fig. 5d**). The position of each band is determined by fitting the peak of the intensity plot using a Gaussian fit and determining the position of the peak maxima.

### Script for intensity profile alignment

```
# Find the best shift using cross-correlation
def find_best_shift(profile1, profile2):
    correlation = correlate(profile1 - np.mean(profile1), profile2 - np.mean(profile2), mode='full')
    shift = np.argmax(correlation) - (len(profile2) - 1)
    return shift

# Align profiles
def align_profiles(profile1, profile2):
    profile2_flipped = np.flip(profile2)  # 180-degree rotation

    # Compute best shifts for both orientations
    shift_0 = find_best_shift(profile1, profile2)
    shift_180 = find_best_shift(profile1, profile2_flipped)

    # Choose the best alignment based on highest correlation
    best_rotation = 0 if abs(shift_0) >= abs(shift_180) else 180
    best_shift = shift_0 if best_rotation == 0 else shift_180

    return best_rotation, best_shift
```

**Table S4.** Position of the sub-bands

| Name sub-band | Control | | | CsA | | | | |
|---|---|---|---|---|---|---|---|---|
| | Position (nm) | Standard deviation (nm) | n | Position (nm) | Standard deviation | n | Difference (nm) | p value |
| b2 | 8.59 | 0.31 | 16 | 8.47 | 0.68 | 13 | 0.12 | 0.552 |
| b1 | 15.95 | 0.40 | 15 | 15.04 | 0.50 | 13 | 0.92 | 2.30E-05 |
| a4 | 21.61 | 0.42 | 7 | 21.39 | 0.29 | 7 | 0.21 | 0.329 |
| a3 | 25.03 | 0.55 | 15 | 25.04 | 0.30 | 8 | -0.01 | 0.952 |
| a2 | 28.63 | 0.89 | 4 | 27.88 | 0.24 | 3 | 0.75 | 0.283 |
| a1 | 32.36 | 0.48 | 15 | 31.78 | 0.38 | 9 | 0.58 | 0.007 |
| e2 | 38.30 | 0.43 | 18 | 37.86 | 0.46 | 13 | 0.44 | 0.013 |
| e1 | 44.57 | 0.68 | 18 | 44.38 | 1.02 | 12 | 0.19 | 0.559 |
| d | 52.94 | 0.44 | 19 | 52.34 | 0.56 | 13 | 0.60 | 0.002 |
| c | 62.50 | 0.55 | 16 | 63.30 | 1.45 | 11 | -0.80 | 0.063 |
| D-band | 67.13 | 0.31 | 15 | 67.11 | 0.45 | 11 | 0.02 | 0.880 |
